# Protocells reproduce morphological and chemical signatures of early microfossils

**DOI:** 10.64898/2026.09.28.753251

**Authors:** Louise Gillet de Chalonge, Kaitlin Field, Róisín Dignam, Maxime Coutant, Corentin Loron, Mark A. van Zuilen, Seán F. Jordan

**Affiliations:** Life Sciences Institute, School of Chemical Sciences, Dublin City University, Glasnevin, Dublin 9, Ireland; School of Natural Sciences, Botany, Trinity College Dublin, Dublin, Ireland; Naturalis Biodiversity Centre, Leiden, 2333CR, The Netherlands

**Author notes:** **Corresponding author:** Seán F. Jordan.

**Keywords:** origin of life, protocells, lipid vesicles, microfossils, biomorphs, biogenicity criteria, biosignatures, protobiosignatures

## Abstract

Identification of early microfossils relies on morphological and chemical criteria assumed to be unique to life^1–4^, but whether abiotic microstructures can satisfy these criteria has not been rigorously tested. Hydrothermal environments preserve microfossils but are also efficient geochemical reactors for abiotic reactions^5^. Protocells form in hydrothermal conditions and represent a plausible abiotic analogue to early microfossils^6^. Here we show that protocells form from simple organic molecules in low-temperature, alkaline hydrothermal conditions in the presence of silica and produce spheres, filaments, and lenticular structures that strongly resemble the morphology of purported microfossils in Archaean silica cherts. FTIR spectroscopy provides overlapping chemical signatures across protocells, microorganisms, and geologic microstructures. In addition, the FTIR R_3/2_ ratio, a chemometric index often used to discriminate domains and determine biogenicity^7–9^, identifies eight different protocell systems as either bacterial, archaeal, or eukaryotic. These data suggest that protocells preserved in ancient hydrothermal deposits can satisfy current biogenicity criteria, undermining the reliability of interpretations of the early fossil record. As life detection is the primary objective for current space exploration endeavours, robust biogenicity protocols that account for abiotic signatures are essential.

## Introduction

Organic microfossils have been reported in many Archaean silica deposits^1–3^ and are suggested to be some of the oldest traces of life on Earth. Some of these organic traces occur in seafloor hydrothermal silica deposits, triggering the discussion as to whether they represent 1) marine microorganisms that lived in the upper water column and later introduced in these seafloor systems by circulating seawater^10^, 2) chemolithoautotrophic microorganisms that lived in such seafloor hydrothermal settings^11^, or 3) abiogenic organic materials directly resulting from hydrothermal reactions with the (ultra)mafic ocean floor^12^. In hydrothermal environments, heated water circulating through rocks dissolves silicate minerals, leading to a high concentration of silica in resulting fluids. As fluids cool down, acidify or go through wet-dry cycles, they oversaturate in silica, leading to the silicification of organic structures^13,14^. This process produces resilient microstructures which can sometimes preserve through diagenesis and metamorphosis over billions of years^1–3^. However, determining whether these microstructures represent true microfossils of early organisms is often controversial, since metamorphism has altered the original organic molecules to macromolecular kerogen, making it difficult to distinguish between abiologic and biologic sources.

In recent years, discoveries of non-living chemical reactions that can produce morphologies remarkably similar in shape and size to ancient microfossils, so-called biomorphs, have reinforced the need to improve our interpretation of geological microstructures^6,15–19^. Beyond morphology, additional biogenicity criteria are used to assess the biological origin of a structure in the rock record in an attempt to overcome potential confusion with abiotic structures^4^. Some commonly used criteria include colonial or community-like behaviour, preferred orientation (i.e., microorganisms forming a biofilm in a specific direction), isotopic fractionation, and spectroscopic ratios^20,21^. However, many of these criteria could be met by abiotic chemical reactions^6,22,23^.

Silica-rich hydrothermal environments have excellent microfossil preservation potential and are also seen as the most plausible settings for the emergence of life. Acting as geochemical reactors, they provide organic molecules that undergo self-assembly and subsequent reaction pathways eventually leading to the first living cells^24^. Along this trajectory, abiotic organic microstructures of varying degrees of complexity are formed including protocell membranes — precursors to biological cell membranes. Recently, it has been shown that protocells form structures that meet some of the morphological and morphometric biogenicity criteria applied to early microfossils^6,25^. We can suppose that, as prebiotic chemistry developed towards life, the distinction between prebiotic systems and living cells becomes even more challenging to discriminate, both on a morphological and molecular level.

To date, most biomorph experiments have focused on inorganic structures^15,16^. There are several reports of potential organic biomorphs^17,18,26^ but the prebiotic plausibility of some of these products remains unclear. There is currently no data on the potential preservation of plausible protocells, the morphological diversity of the resulting structures, and whether the structures could be distinguished morphologically and chemically from fossilised early cells. This presents a major challenge for understanding the origin and evolution of early life on Earth — this challenge is arguably even greater when attempting to detect life on other planetary bodies such as Mars. Mars could have fostered life early after its formation when conditions were similar to those of early Earth. The remains of early Martian life, therefore, could be preserved in the Noachian rock record. However, life may also never have arisen on Mars, potentially leaving behind a record consisting solely of abiotic and/or prebiotic organic signatures^27^. Interpreting this rock record without understanding the preservation potential and diagnostic signatures of abiotic microstructures is a daunting task, one that should be tackled prior to Mars sample return.

Here, we investigated the self-assembly of protocells in silica and compared them with data from extant microorganisms and Archaean putative microfossils. Protocells were formed from decanoic acid (DA) and 1-decanol (DOH) in analogue hydrothermal vent conditions (70 °C, pH 6 to 13) in the presence of silica (Na_2_SiO_3_; 300 ppm Si). These conditions focus specifically on simulating a seafloor hydrothermal setting with low-temperature, neutral to alkaline fluids, although it has been shown that protocells can form in a wide range of pH, salinity and temperature^28–31^. Protocells remained stable in solution at pH 9 for up to 48 hrs at 70 °C and for several weeks at ambient temperature. We observed that DADOH protocells formed in silica produced a variety of morphologies including isolated and clusters of spheres, filaments, and lenticular structures of size and shape consistent with microfossils. A total of eight protocell compositions formed in silica (C_10_-C_15_ fatty acid and 1- alkanol pairs; mix of C_10_-C_16_ fatty acids and alkanols; and a pair of C_10_ fatty acid and isoprenoid alcohol) were analysed by Fourier transform infrared (FTIR) spectroscopy. Resulting spectra overlapped with those from Archaean putative microfossils. In addition, the R_3/2_ ratio — a proxy commonly applied to microfossil FTIR data to determine biogenicity and discriminate between bacterial, archaeal, and eukaryotic sources — places protocells across all domains of life with values that match those of Archaean microfossils^8,32^. These data further confirm that this ratio is insufficient to accurately interpret lipid signatures in rock samples^9,33^. Our results highlight the need to advance our understanding of the preservation potential of abiotic and prebiotic organic signatures to improve our interpretation of Archaean and extraterrestrial samples.

## Results

### Protocells form in the presence of silica

To assess protocell formation in silica-rich environments, protocells were formed with DA and DOH across a pH range from 6 to 13, in deionised H_2_O only or in Na_2_SiO_3_ solution (300 ppm Si). Optical density measurements at 480 nm reflect the state of fatty acids in solution, as different structures (micelles, vesicles or droplets) exhibit different light-scattering properties. Protocells self-assembled across the entire pH range, in the presence and absence of silica (Fig. 1a), confirmed with UV-Vis spectroscopy and fluorescence microscopy. The surface charge of protocells was measured with dynamic light scattering (DLS) across a pH range from 7 to 14 (Fig. S1). Protocells in both control conditions and exposed to silica have the same surface charge across the range, -75 mV from pH 7 to 13. The FTIR spectra of DADOH protocells in control conditions and in the presence of silica show peaks at 2960, 2925, 2855, 1544, 1460 and 1410 cm^-1^ (Fig. 1b) which correspond to C-H bonds in aliphatic chains and are common diagnostic peaks in microfossils interpretation.

**Figure 1:**
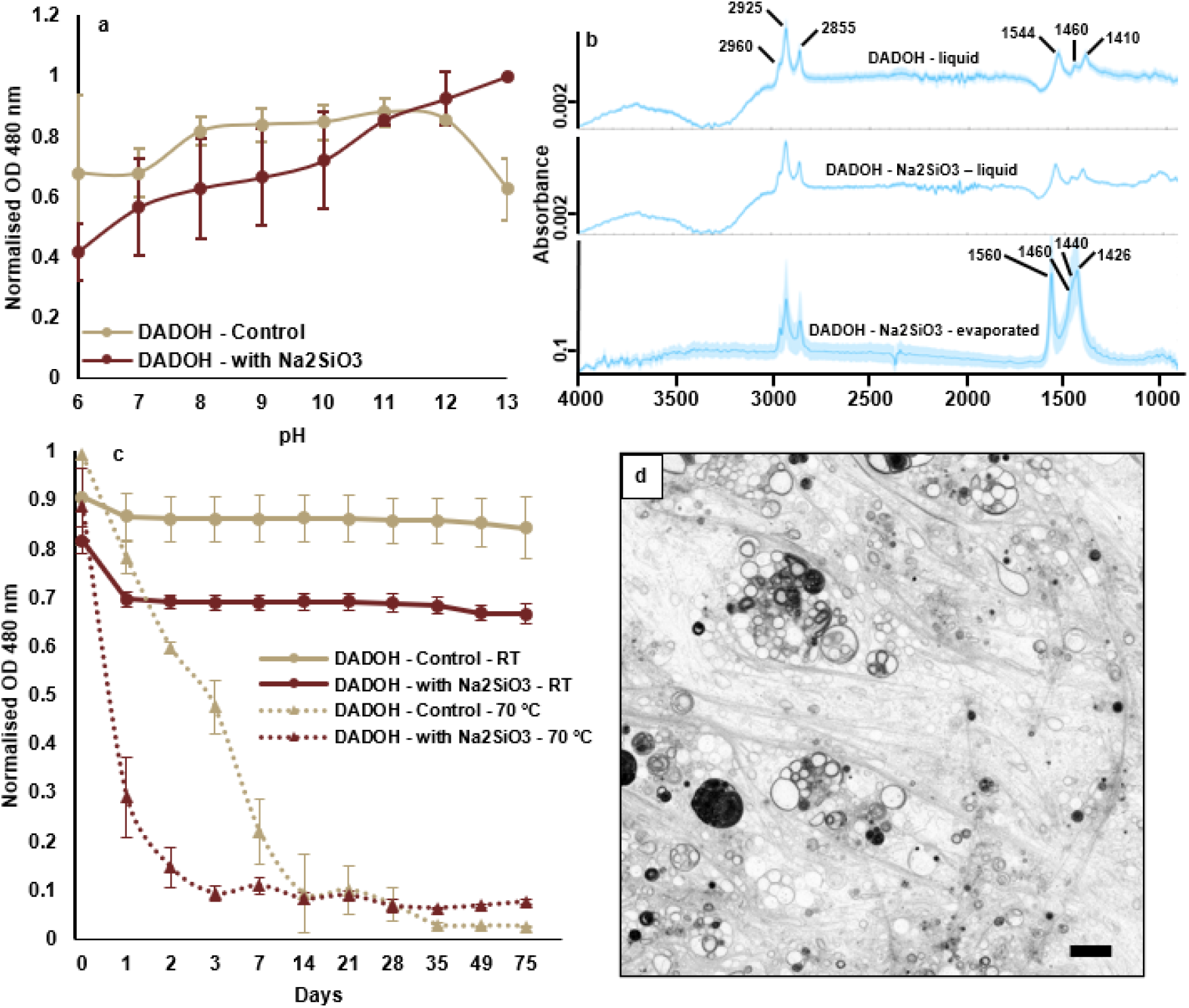
Protocells can self-assemble in the presence of silica. (a) Absorbance measurements at 480 nm of DADOH protocells across a pH range from 6 to 13, in control conditions (yellow) or exposed to Na_2_SiO_3_ (300 ppm Si) (dark red). (b) FTIR spectra of DADOH protocells in liquid solution, in liquid solution with Na_2_SiO_3_ (300 ppm Si), or prepared in solution with Na_2_SiO_3_ and evaporated. (c) Absorbance measurements at 480 nm, showing the stability over 8 weeks of DADOH protocells at pH 9, stored at room temperature (circles) or at 70 °C (triangles), in control conditions (yellow) or exposed to 300 ppm Si (dark red). (d) CSLM picture of DADOH protocells formed in the presence of 300 ppm Si and stained with Rhodamine 6g. Scale bar = 10 µm.

Protocells were stable in solution for up to 8 weeks at room temperature at pH 9, with and without silica (Fig. 1c). At 70 °C, the optical density for protocells without silica decreased consistently over 7 days. In samples exposed to silica, protocells disappeared quickly over 2 days likely forming a mixture of dissolved monomers and micelles. Experiments were repeated with colloidal silica instead of Na_2_SiO_3_ at 300 ppm Si and results were consistent (Fig. S2). Additionally, protocells were prepared in a highly concentrated silica mixture containing both colloidal silica and Na_2_SiO_3_, with a final concentration of 0.5 M silica, to simulate highly concentrated fluid (Fig. S3). The silica mixture was brought to neutral pH with addition of HCl, forming a gel after a few minutes at room temperature. This simulates a silicification process that can happen naturally in hydrothermal environments or silica sinters. The pH of the Archaean ocean was mildly acidic (ca. 6.5-7), which was primarily due to an atmosphere rich in CO_2_, and temperature estimates range between 30 and 70 °C^34–36^.

Warm, silica-rich fluids, when mixed with colder or more acidic ocean waters, would precipitate and form a silica gel. Here, the presence of protocells in the silica mixture before gelation was confirmed with confocal laser scanning microscopy (CSLM) (Fig. S3a). Protocells were less abundant than in samples with lower silica concentrations, but still present and visible. Following gelation, large protocells were still observable in fluid inclusions within the gel with optical microscopy (Fig. S3b).

These results show that protocells can form in conditions relevant to prebiotic hydrothermal environments, can persist in the presence in silica, and can be entombed in a silica gel, demonstrating that protocells could persist during natural silicification processes in Archaean hydrothermal vents.

### Protocell morphologies are comparable to Archaean microstructures

The formation of protocells in different experimental conditions (in the presence or absence of silica; in solution or evaporated) led to a variety of morphologies that are strikingly similar to microstructures found in Archaean cherts, as observed under CSLM and scanning electron microscopy (SEM). Protocells in control conditions formed spheres, free-floating in solution or emerging from droplets (Fig. 2a). In the presence of silica, protocells formed more complex aggregates with spheres, filaments and multivesicular structures (Fig. 2b). After evaporation, we observed filaments (Fig. 2c) as well as spheres, although the carbonaceous nature of these spheres has not been confirmed (Fig. 2d). Across different experimental conditions, we were able to abiotically mimic most Archaean morphologies: spheres (Figs. 3a to 3e), aggregates of spheres (Figs. 3f to 3m), dividing or coalescing spheres (Figs 3n to 3r), lenticular structures (Figs 3s to 3v), filaments (Figs 3w to 3z) and chains (Figs 3α to 3γ). Most commonly, protocells in solution form spheres of varying sizes (ca. 0.5 to 20 µm) (Figs. 2a and 3d). Carbonaceous spheres are one of the most common morphologies reported for microfossils. For example, a population of spheres with a size distribution of 4 to 22 µm has been described in chert from the 3.0 Ga Cleaverville Formation, Pilbara craton, Western Australia^37^ (Fig. 3c). The biogenic origin of these spheres is not yet well established. Protocells formed in solution with silica were more likely to form multivesicular structures (Figs. 2b and 3e), with one large vesicle encapsulating one or several smaller vesicles, which has also been described in cherts from the SPF^38^. The presence of silica led to the formation of clusters of spheres as well as complex aggregates composed of spheres and filaments (Figs. 3h-j and 3m). Clusters of carbonaceous spheroids of 20 to 40 µm with a foam-like envelope were described in the SPF (3.4 Ga) and were interpretated as biological^39^ (Fig. 3l). In samples consisting of protocells formed in the presence of silica and evaporated, we observed spheres with a diameter of 1.22 µm ± 0.23 (measured on 50 specimens) present in a colony-like group (Fig. 3m), similar to spheroids and globules found in the SPF^39,40^ (Fig. 3k). Protocells will sometimes spontaneously divide in solution, leading to pairs or aggregates of coalescent spheres (Figs. 3q and 3r), a morphology that has also been seen in the Cleaverville Formation^37^ as well as in tephra fragments^41^ and abiotic carbonaceous spheroids^38^ from the 3.4 Ga Strelley Pool Formation (SPF) (Figs. 3n to 3p).

**Figure 2:**
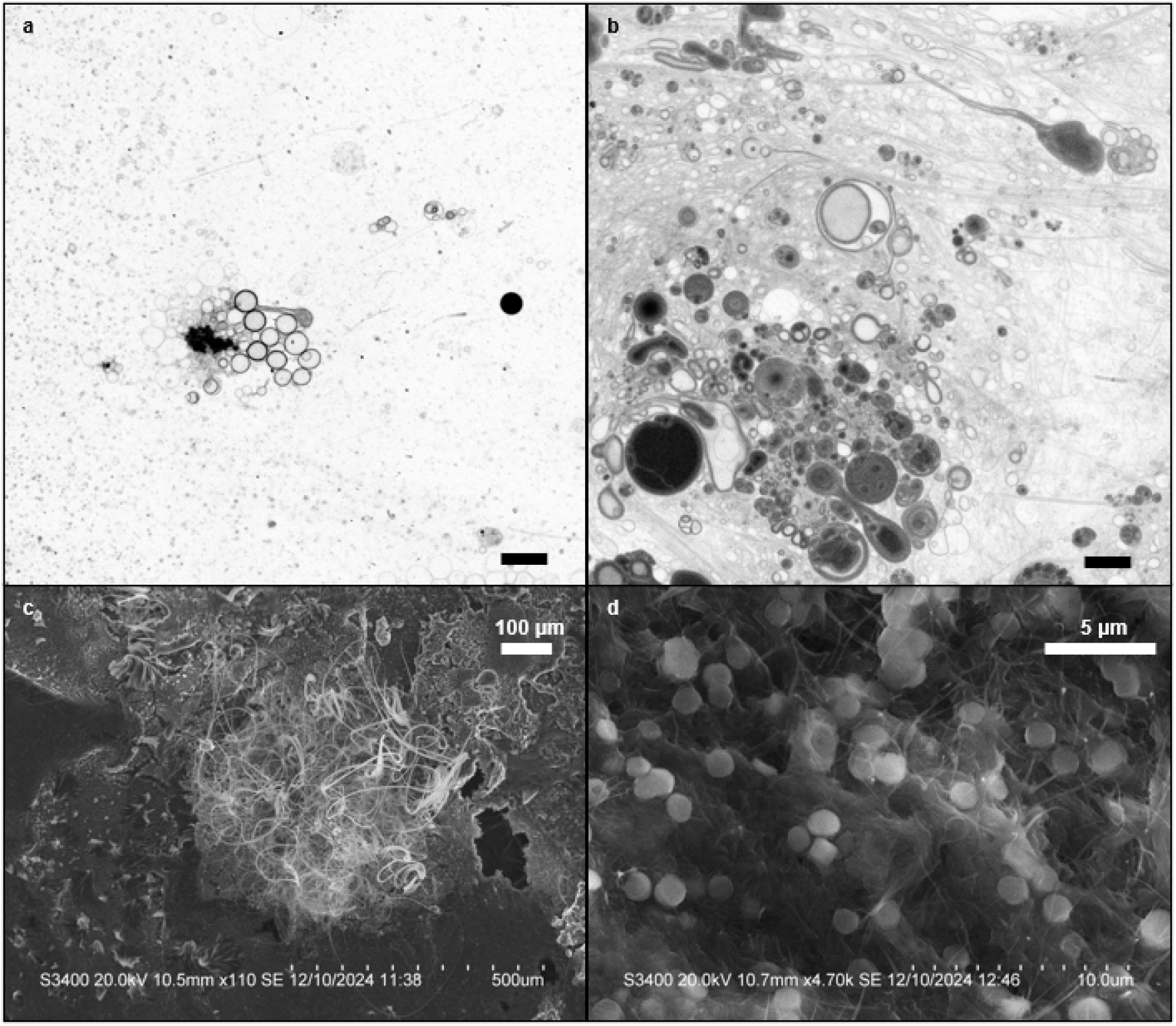
CSLM and electron microscopy of DADOH protocells. (a-b) CSLM micrographs of protocells (a) in control conditions and (b) formed in the presence of 300 ppm Si, stained with Rhodamine 6g. Scale bar = 10 µm. (c-d) Scanning Electron Microscopy (SEM) images of DADOH protocells formed in solution with 300 ppm Si then evaporated, showing different types of structures, such as (c) filaments and (d) spheres present in a colony-like group.

**Figure 3:**
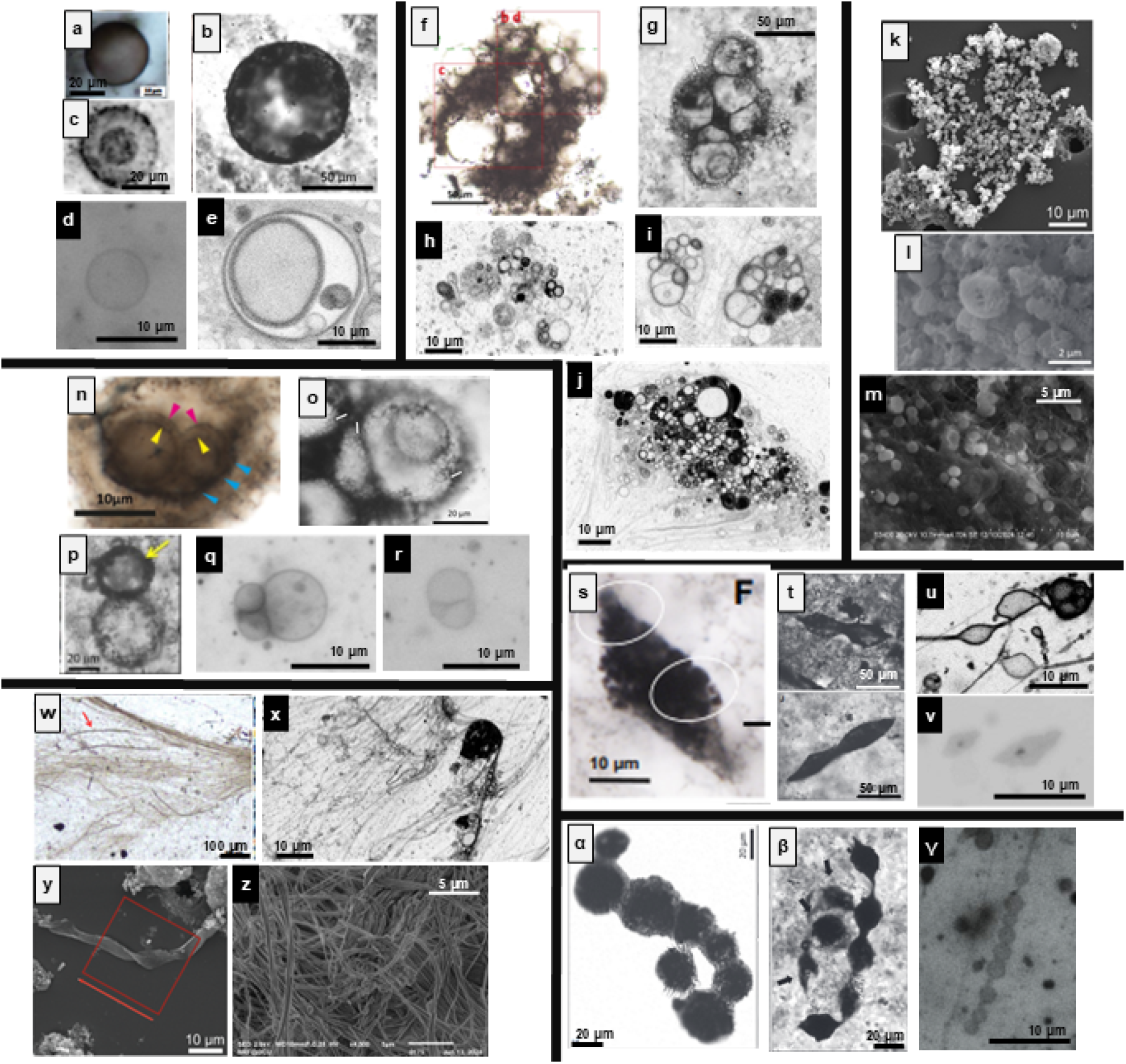
Comparison of carbonaceous structures found in putative Archaean microfossils and morphologies of protocells formed in the context of this study. Lettering in white font over black background indicates biomorphs produced in this study, while lettering in black font over white background are published purported microfossils. (a-e) Spheroidal structures. a. Sphere in black chert from the 2.4 Ga Turee Creek Group, Western Australia.^42^ b. Hollow sphere from the 3.4 Ga Waterfall Locality, Strelley Pool Formation (SPF), Western Australia.^39^ c. Spheroidal microstructure from the 3.0 Ga chert of the Cleaverville Formation, Western Australia.^37^ d. DADOH protocells formed and kept for 7 days at room temperature in the presence of 300 ppm Si, stained with Rhodamine 6g, optical microscopy. e. DADOH protocells formed in the presence of 300 ppm Si, picture taken after preparation, stained with Rhodamine 6g, CSLM. **(f-m) Aggregates of spheroidal structures** f. Cluster of spheroids from the 3.4 Ga SPF, Western Australia.^38^ g. Clustered hollow spheres of similar sizes, described as being surrounded by a foam-like texture, from the 3.4 Ga SPF, Western Australia.^39^ h-j. Aggregates of protocells formed in the presence of 300 ppm Si, picture taken after preparation, stained with Rhodamine 6g, CSLM. k. SEM picture of acid-macerated spheroids from the 3.4 Ga SPF, Western Australia.^40^ l. SEM picture of acid-macerated carbonaceous globules from the 3.4 Ga SPF, Western Australia.^39^ m. SEM pictures of spheres found in a DADOH protocell sample, formed in solution with silica and evaporated at room temperature. **(n-r) Division-like patterns** n. Sphere encapsulating multiple spheroidal vesicles, from the 3.4 Ga SPF, Western Australia.^38^ o. Spheres from the 3.4 Ga SPF, Western Australia.^39^ p. Spheres found in tephra fragments from the 3.4 Ga SPF, Western Australia. ^41^ q-r. DADOH protocells formed and kept for 7 days at room temperature in the presence of 300 ppm Si, stained with Rhodamine 6g, optical microscopy. **(s-v) Lenticular structures** s-t. Lenses found in chert in the 3.4 Ga SPF, Western Australia.^3,39^ u-v. Lenticular structures formed in DADOH protocells exposed to 300 ppm Si (respectively, CSLM and optical microscopy). **(w-z) Filaments** w. Filamentous structures, described as “rivers of straw-like filaments”, found in the 2.4 Ga Turee Creek Group, Western Australia.^42^ x. Similar rivers of organic filaments, very common in protocells solution, especially when the sample is starting to dry. DADOH solution formed in the presence of 300 ppm Si, stained with Rhodamine 6g, CSLM. y. SEM picture of acid-macerated isolated filament found in chert in the 3.4 Ga SPF, Western Australia.^40^ z. SEM picture of filaments found in a DADOH protocells sample, formed in solution with silica and evaporated at room temperature. **(α-γ)) Chains** **α-β**. Acid-macerated and silica-embedded chains of lenses found in the.4 Ga SPF, Western Australia.^39^ **γ** . Chain of DADOH protocells prepared in the presence of 300 ppm Si, stained with Rhodamine 6g, optical microscopy. Chains sometimes form when protocells are dividing and are attached to the slide on one end but are being drawn by the flow in solution.

Filamentous structures, described as “river of straw-like filaments”, are shown in Paleoproterozoic samples from the Turee Creek Group, Western Australia (2.4 Ga)^42^ and are similar to filaments found in protocell solutions (Figs. 3w and 3x). In our study, organic filaments of ca. 2 µm of diameter were almost ubiquitous in solution but also in samples that were dried at 50 °C and imaged with SEM, and isolated filaments of similar size were found in the SPF^40^ (Figs. 3y and 3z). We have observed lenticular structures forming in solution or on evaporated slides, with morphologies similar to lenses that have been previously described in the SPF cherts^3,38,39^ and often interpreted as biological (Figs. 3s to 3v). Chains of carbonaceous lenses were observed in 3.4 Ga SPF chert with significant size variation between the different units, on average 20 µm (Figs. 3α and 3β). In solution, protocells sometimes form chains, especially when one part of a vesicle is attached to a surface (e.g., an artefact of the microscopy slide) and the vesicles start dividing while being drawn by the flow in solution (Fig. 3γ).

### FTIR spectroscopy does not distinguish between protocells, microorganisms or Archaean microfossils

In addition to morphology, we evaluated whether FTIR spectroscopy could discriminate abiotic protocells from living cells and microfossils. We examined a range of prebiotically plausible membrane compositions (C_10_ to C_15_ fatty acid and alkanol pairs, C_10_ to C_16_ fatty acid and alkanol mix, C_10_ fatty acid and isoprenoid alkanol pair), some of which have already been shown to mimic microfossil morphologies^6^. FTIR analysis was conducted immediately after preparation of protocells in solution, and after a week at room temperature or 70 °C, in the absence or presence of silica. Additionally, micro-FTIR was conducted on C_10_, C_13_ and C_15_ fatty acid/1-alkanol pair protocells evaporated on BaF_2_ slides.

The FTIR spectra of all protocells in solution (in the absence or presence of silica) and protocells evaporated on BaF_2_ slides show similar profiles. The full spectra of DADOH protocells are shown as an example (Fig. 1b). The main peaks of interest correspond to C-H bonds in aliphatic chains. In the region 2700-3000 cm^-1^, 3 peaks were identified in samples in solution and evaporated samples: 2855 cm^-1^ (CH_2_ symmetric stretch), 2925 cm^-1^ (CH_2_ asymmetric stretch), and 2955-2960 cm^-1^ (CH_3_ asymmetric stretch)^43^. These 3 peaks are often discussed in microfossil analysis and used to support a biological origin for a microstructure, for example in 3.0-3.4 Ga carbonaceous chert from Western Australia described in Sugitani et al., 2026 and Igisu et al., 2018. To further characterise this region, we calculated the R_3/2_ ratio for all samples. Results show that the R_3/2_ of all abiotic protocells are in the range of 0.15 to 0.85 (Fig. 4), with values increasing with chain length. This aligns with predicted values calculated using the number of CH_3_ and CH_2_ moieties within each molecule. Micro-FTIR results are consistent with bulk FTIR in liquid solutions. Most importantly, the ratios also fall in the range of microfossils characterised in Sugitani et al., 2026 and Igisu et al., 2018. Moreover, they were consistent with R_3/2_ values of whole cells of extant archaeal, bacterial and eukaryotic microorganisms^7^. Deconvolution of the aliphatic interval is sometimes conducted to analyse this region of the spectrum, using 5 main C-H contributions (CH_3_ and CH_2_ asymmetrical stretching; CH_3_ and CH_2_ symmetrical stretching; and CH stretching). Alternative processing of FTIR data can yield varying R_3/2_ values based on the type of preprocessing (for example, values ranged mostly between 0.1 and 0.4 for fatty acid-alkanol pairs if the spectra was deconvoluted before calculating the R_3/2_), which could potentially lead to incorrect interpretation (Fig S4). These findings highlight significant limitations in the use of R_3/2_ ratios for the interpretation of ancient microstructures.

**Figure 4:**
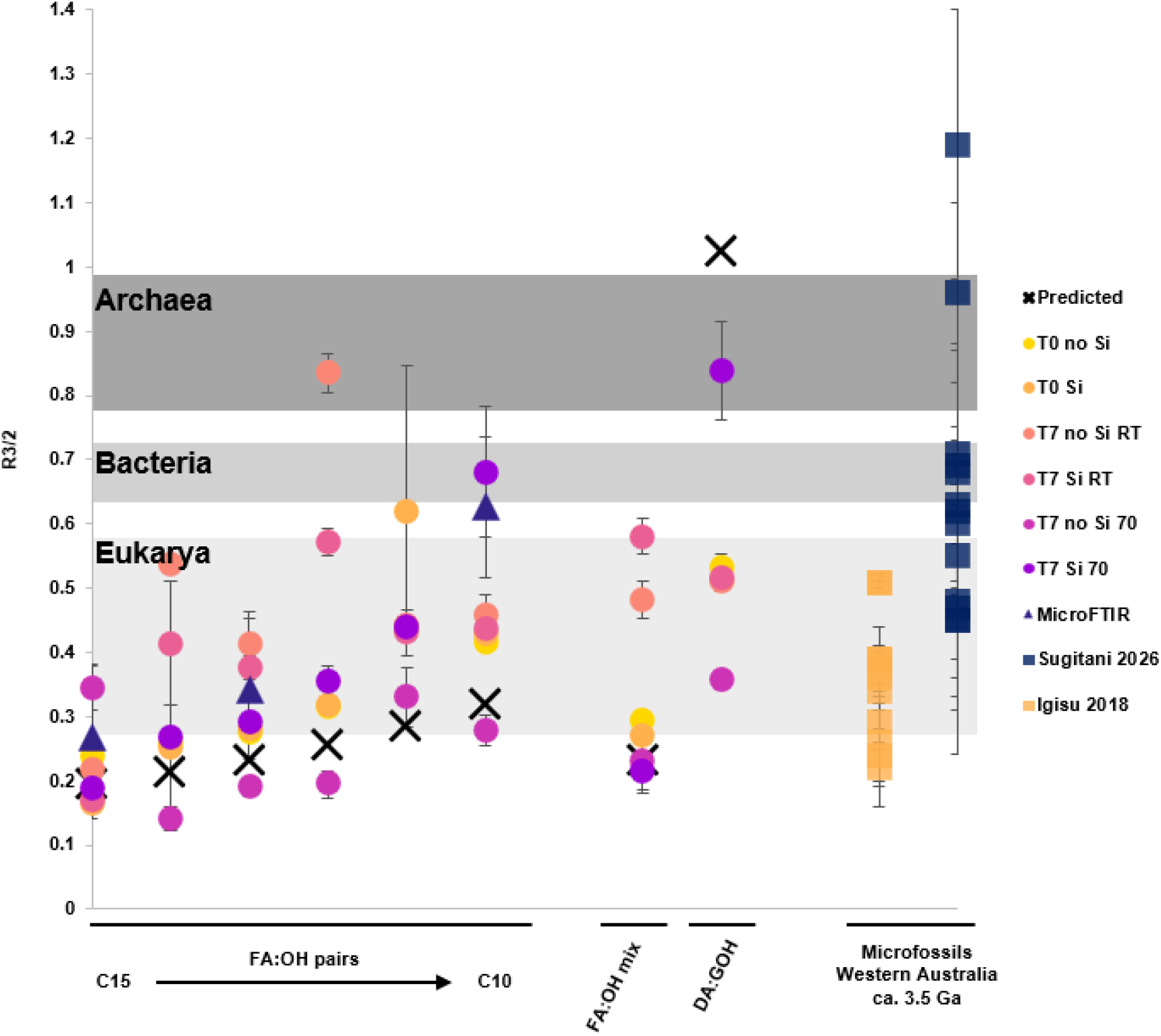
The R_3/2_ values of protocells fall in the same range as biotic samples as well as purported Archaean microfossils. Comparison of the R_3/2_ value of protocells in solution (circles), evaporated protocells (triangle), putative Archaean microfossils (squares), and whole cells from extant living organisms (range in greyscale), all calculated from FTIR spectra, as well as predicted values for protocells (cross). FTIR spectra of protocells in solution were taken right after preparation (T0) or after 7 days (T7) at 70 °C or at room temperature, in control conditions (no Si) or in the presence of 300 ppm Si. Micro-FTIR spectra were taken of protocells evaporated right after preparation. Values from Igisu et al., 2018 were measured in microstructures found in carbonaceous clots in ca. 3.5 Ga hydrothermal vein deposits from the Dresser Formation, Western Australia. Microfossils from Sugitani et al., 2026 are organic-walled spheroid and ellipsoid microfossils from the ca. 3.0 Ga Farrel Quartzite in Pilbara Craton, Western Australia. The background range in greyscale represents the measured R_3/2_ values for whole cells of extant organisms (including eukaryotes, bacteria and archaea) in Igisu et al., 2009. Predicted R_3/2_ values of protocells were calculated following the correlation in Igisu et al., 2009 (R_3/2_ = 2.56 × [CH_3_/CH_2_], r2 = 0.64), after calculating the amount of CH_3_ and CH_2_ in each amphiphilic molecule used.

In the region 1000-1500 cm^-1^, peaks differ between samples in solution and evaporated samples. For samples in solution, peaks were found at 1544 cm^-1^ (C=C stretching or COOH/COO^-^), 1460 cm^-1^ (CH_x_ bend or -COH functions), 1410 cm^-1^ (CH bend or OH)^43,44^. The micro-FTIR of evaporated spectra contains peaks at 1560 cm^-1^, 1460 cm^-1^, 1440 cm^-1^, and 1426 cm^-1^, which can correspond to carboxyl functional groups as well as different bending of C-H bonds^43^. Peaks around 1550-1560 cm^-1^ have also been described in extracted microfossils from Sugitani *et al*., 2026. However, most organic peaks in the 1000- 1500 cm^-1^ region are overlooked in microfossil studies as they are often covered by the peaks of Si-O bonds from the silica matrix.

If protocells were entombed in a silica matrix that underwent diagenesis through geological time, the resulting spectra could be mistaken for having a biological origin, with key peaks in the 2700-3000 cm^-1^ region being undistinguishable from those of fossilised cell membrane lipids, and organic signatures in the 1000-1500 cm^-1^ region being uninterpretable due to the overlapping with intense peaks of SiO_2_ bonds.

### Abiotic organic structures may be indistinguishable from Archaean microfossils

Our results show that protocells form and persist in hydrothermal conditions in the presence of silica and reproduce some of the morphological and chemical signatures commonly used to interpret the biogenicity of early microfossils. The addition of 300 ppm Si (ca. 10 mM) did not affect the self-assembly of protocells composed of amphiphilic molecules known to be produced by hydrothermal Fischer-Tropsch type synthesis^5^. Protocells formed across a wide range of pH and at temperatures relevant to hydrothermal environments. In modern alkaline hydrothermal systems, the concentration of silica in hydrothermal fluids ranges from 0.001 to 1.5 mM^45,46^. However, geochemical analysis of Archaean cherts suggests that the Archaean ocean had high concentrations of silica (ca. 1.9 mM)^14,47^ and silica concentrations would have been higher than 1.9 mM in hydrothermal fluid. We show here that the presence of silica, even in concentrations up to 500 mM, would not have been an obstacle against the formation and possible evolution of protocells in these environments. Furthermore, protocells were stable at ambient temperature for up to 8 weeks but transitioned to micelles and monomers in 1-2 days at 70 °C. Hydrothermal environments exhibit pH and temperature gradients, with warm, alkaline fluids near the vent source transitioning to colder, slightly acidic conditions in the surrounding ocean. This gradient could favour the emergence and evolution of protocells. Bilayer vesicles could quickly form, disintegrate and re-form in the warm hydrothermal pores, allowing for increased mixing of membrane lipids and protocell contents. This would provide for a wider chemical reaction space which may have been important for early prebiotic chemistry. More developed protocells could then achieve increased stability upon incorporation of phospholipids or when transported to colder, more acidic ocean waters, which would favour long-term stability and colonisation of surrounding environments.

Once formed, protocells could persist in a highly concentrated silica solution (0.5 M) and after gelation. Silica gels were likely pervasive in the Paleoarchaean ocean, especially surrounding hydrothermal vents^48^. Although the rate of gel formation depends on physicochemical conditions (temperature, pH, silica concentration, ionic strength), it would range from a few days near the hydrothermal source to a couple of weeks or months further away^49^. Protocells might persist long enough in these silica gels to be preserved during silicification processes.

Protocells formed in the presence of silica produced a diversity of morphologies that are strikingly reminiscent of carbonaceous structures found in Archaean cherts. This includes spheres, clustered spheres, chains of spheres, filaments, and lenticular structures, that parallel reported shapes in Archaean samples, mainly in the Strelley Pool Formation (3.4 Ga)^3,37–42^. Although the main morphology of protocells in the absence of silica is spheroidal, protocells assembled in the silica solutions formed more complex structures including aggregates of spheres and filaments and multilamellar and multivesicular compartments.

Minerals, including silica, have been shown to influence the formation and aggregation of protocells, mainly depending on surface charge and isoelectric point^50^. Free silanol groups in solution could also interact with amphiphiles and potentially bind to protocell membranes, further influencing morphology and aggregation. Division patterns can result from protocell fission and division which happen spontaneously in solution^51^. The formation of filaments may be enhanced by evaporation or by the presence of salts^31^. Overall, physico-chemical conditions occurring naturally in hydrothermal environments, such as the presence of salts, mineral surfaces, or evaporation cycles, could lead to the diversity of morphologies observed in this study. This demonstrates that morphological complexity is not necessarily linked to a biotic origin in Archaean samples, including colonial or community-like clusters of carbonaceous spheres and lenticular structures, which have long been considered biogenicity criteria^20^. The null hypothesis should be an entirely abiotic system when interpreting these types of structures based on morphological observations, and more work is needed to provide a complete abiotic baseline for early Earth environments, including deeper knowledge of depositional and taphonomic conditions.

In addition to morphology, we observed that chemical data obtained by FTIR could not be used as reliable biosignatures. We demonstrated that the R_3/2_ ratio could not discriminate between abiotic protocells and biological membranes in samples derived from hydrothermal silica-rich environments. The R_3/2_ values of protocells, formed in different conditions, overlap with values reported for extant cells within the three domains of life^7^ and with values measured on putative microfossils from Archaean cherts^8,32^. We also observed that variations in preprocessing lead to different R_3/2_ values, highlighting the potential for methodological bias when using this ratio for sample interpretation. As spectroscopic techniques more broadly, and FTIR specifically, are primary techniques for life detection missions in our solar system, caution should be used when interpreting these data especially considering the lack of data for potentially preserved abiotic signatures.

## Conclusion

We conclude that protocells formed by simple organic molecules could have self-assembled in the presence of silica in Archaean hydrothermal environments. Although the high temperatures of hydrothermal fluids could affect the stability of the protocells over time, they could have emerged and remained stable when increasing in complexity or mixing with the colder waters of the ocean. Protocells could be entombed in silica rich matrices and remain preserved in Archean cherts. The morphologies resulting from the formation of protocells in the presence of silica mimic most previously reported microfossil morphologies from Archean cherts, eliminating morphology as a reliable biosignature. Finally, the associated chemical signatures of preserved protocells are indistinguishable from specimens with a proposed biological origin in Archaean cherts using FTIR spectroscopy. This work highlights the importance of considering the abiotic and prebiotic baseline for life detection on the early Earth and in the context of space exploration.

## Material and methods

## Materials

All membrane-forming molecules (C_10_ to C_16_ fatty acids, C_10_ to C_16_ alkanols, geraniol) were purchased from Sigma-Aldrich.

Two different sources of silica were chosen and used for all experiments: a sodium silicate solution (Na_2_SiO_3_, 27 wt% SiO_2_ and 10 wt% NaOH) and colloidal silica (LUDOX HS-40).

Both solutions have been commonly used in previous artificial silicification studies and present different properties, including the size of silica particles. LUDOX HS-40 and Na_2_SiO_3_ were purchased from Sigma-Aldrich. Rhodamine 6g was used for imaging protocells in fluorescence and CSLM.

Samples for absorbance, microscopy and Dynamic Light Scattering were prepared in Falcon tubes. For FTIR analysis, experiments were conducted in organic-free conditions, in glassware furnaced at 550 °C for 12 hours. Spatula, tweezers, or any additional tool used for the experiments were cleaned with organic solvents. Before use, stock solutions of reagents (NaOH, HCl, Rhodamine 6g, silica solutions) were filtered with a syringe and a 0.22 µm filter pore membrane to remove any particle or possible bacterial contamination.

### Preparation of protocells

Protocells were always prepared fresh before each experiment, following the standard protocol from Monnard and Deamer^52^ (2022) to form bilayer vesicles from amphiphilic molecules. Solutions were prepared in a heatblock (Fischerbrand Isotemp) at 70 °C, with all reagents heated at the same temperature to simulate hydrothermal fluid temperature. Decanoic acid (CH_3_(CH_2_)_8_COOH) (DA) and 1-decanol (CH_3_(CH_2_)_9_OH) (DOH) were used for the preparation of protocells in all experiments, unless stated otherwise. Both molecules are relevant for prebiotic environments, have been widely used as models for vesicles and protocell studies, and have previously shown stable formation of vesicles in hydrothermal conditions. Solutions were prepared in deionised water, with an equimolar ratio of DA and DOH. To prepare 7 mL of a 20 mM DADOH 1:1 solution, 25.4 µL of DA was added to 3500 µL of deionised H_2_O and the solution was vortexed. At this stage, the fatty acid is in the state of droplets and the solution is slightly opaque. 3449.1 µL of 0.5 M NaOH was added to increase the pH of the solution (ca. pH 12), leading to the deprotonation of the fatty acid heads and the formation of micelles. Finally, 25.1 µL of DOH was added. The colour of the solution changing from transparent to opaque and foamy after addition of decanol indicates the transition from micelles to vesicles. Protocells of different compositions were prepared following the same protocol, with volumes of amphiphiles, H_2_O and NaOH being adjusted to obtain the right final concentration and pH.

The formation of protocells was consistently confirmed with microscopy observation as well as absorbance measurements at 480 nm on the Shimadzu UV-1800 UV-spectrophotometer. Optical microscopy was conducted on the Nikon Eclipse Ts2R microscope. The pH was measured with a ThermoScientific Orion Star A211.

### Experimental silicification

For the experimental silicification of protocells in aqueous solution, samples containing 20 mM DADOH were prepared in the presence of 300 ppm Si. This concentration of silica was chosen to be in line with previous artificial silicification studies^53,54^. A fresh stock solution of the silica source (Na_2_SiO_3_ or colloidal Si) was prepared at 70 °C in deionised water right before use, at 600 ppm Si. 7 mL of DADOH solution was prepared as described above, with the exception that 3500 µL of H_2_O was replaced by 3500 µL of 600 ppm Si solution, for a final concentration of 300 ppm Si.

For long-term exposure to silica, the pH was set to 9 in all samples to ensure that all solutions have the same alkaline conditions, in line with environmental conditions in hydrothermal environments. Samples were prepared with a final concentration of 300 ppm Si and 40 mM DADOH 1:1, and kept in closed tubes, at 70 °C in a heat block or at room temperature (ca. 20 °C), in the dark, for 8 weeks. The pH, absorbance and FTIR spectra were obtained immediately after the preparation of the solution, followed by weekly measurements. Micrographs were taken at each time point. Every experiment was made in 3 replicates.

Entombment of protocells in a silica gel was prepared following a protocol derived from^55,56^. 1 mL of sodium silicate was gently mixed with 1 mL of colloidal Si, followed by addition of 1 mL of a vesicle solution, for a final concentration of 20 mM DADOH 1:1 and 0.5 M Si. Then, 4 M HCl was added until neutralisation (ca. 80 µL). Gelation occurred within a few minutes at room temperature.

### Optical microscopy

Fluorescence microscopy was conducted on a Nikon Ts2R microscope. Cover glasses (VWR 22x50 mm, Thickness No. 1) were used instead of microscopy slides to facilitate imaging on an inverted microscope. Protocells were observed with the addition of 0.1 mM Rhodamine 6g directly on the slide (with a 1:5 ratio of Rhodamine 6g to vesicle solution) using a 505 nm filter, and 40x or 100x objectives.

For the imaging of protocells in a silica gel, a small amount of gel (ca. 1 mg) was added to a slide and observed on the Nikon Ts2R microscope on brightfield, using 40x and 100x objectives.

CSLM was conducted on the Leica SP8 Confocal STED microscope. Slides were prepared the same way as for fluorescence microscopy, and observe with a laser at 525 nm and a Hyd 1 SMD 1 detector set at 530 to 590 nm, using a 63x objective with immersion oil.

All microscopy images were processed in Fiji for inversion of LUTs, adjustment of brightness and contrast where needed, and addition of a scale bar.

### Scanning Electron Microscopy

Samples were imaged on a Hitachi S3400 scanning electron microscope (SEM). Solutions of protocells (20 mM DADOH 1:1) and protocells with 300 ppm Si, were prepared as described previously, and evaporated directly on a carbon tab, on an Al SEM stub, for 24 hours at room temperature. Samples were imaged on the Hitachi S3400 at 20.0 kV with the secondary electron detector, at a working distance of 10 mm.

### Surface charge measurement with Dynamic Light Scattering

The surface charge of protocells in solution was measured with Dynamic Light Scattering (Malvern Zeta Sizer Ultra), in the complete pH range from 0 to 14. Each experiment was done in triplicates. Solutions of 20 mM DADOH 1:1 were prepared fresh with the pH set by addition of HCl or NaOH. Samples were analysed immediately after preparation in folded capillary zeta cells (Malvern Panalytical DTS1070) to avoid sedimentation. Each sample was measured a minimum of 3 times at 70 °C, with 60 s of equilibration time. A refractive index of 1.45 and absorption of 0.001 were used. For all samples, the dispersant was deionised water, with a refractive index of 1.33 and viscosity of 0.8872 mPa.s at 25 °C.

### Fourier-Transform Infrared spectroscopy

Fourier-Transform Infrared spectra were collected immediately after preparation of protocells in solution (T0), and after a week of storage at room temperature (ca. 20 °C) or 70 °C, in the dark (T7). Spectra were taken on a Perkin Elmer Spectrum Two FT-IR, in the 450 to 4000 cm^-1^ range, with a resolution of 4 cm^-1^. Each spectrum is the average of 4 scans, and there are three replicate per experiment. Initially, a background spectrum of H_2_O was subtracted. However, a residual negative band in the 3500-3000 cm^-1^ region was observed due to O-H bonds of water and NaOH in the samples. A background spectrum of NaOH in H_2_O, or NaOH with 300 ppm Si, at the same temperature as the sample, was taken and subtracted from each sample spectrum. Although this mostly corrected the negative band, some residual band is still present, due to slight differences between the background and samples (e.g., water content, temperature, hydrogen bonds).

For FTIR coupled with microscopy, samples were analysed on a Nicolet raptIR+ FTIR microscope in transmission mode. Spectra were taken on 20 points across the sample evaporated on a BaF_2_ slide, in the 900 to 4000 cm^-1^ range, with a resolution of 8 cm^-1^. Each spectrum is the average of 8 scans. A background spectrum was taken on a clean BaF_2_ slide and subtracted from all spectra.

All spectra were pre-processed on Quasar (version 1.11.1), an extension of Orange software designed for spectroscopy analysis. Spectra were truncated to keep the region from 2750 to 3050 cm^-1^, which contains the relevant peaks for the R_3/2_ analysis. A positive rubberband baseline correction was applied, followed by a Gaussian smoothing of 2. The R_3/2_ was calculated as the ratio between the height of the peak at 2960 and 2925 cm^-1^ (CH_3_ and CH_2_, respectively).

Alternatively, after initial pre-processing, spectra were further processed by applying Savitzky-Golay filter, with a window of 11, a polynomial order of 2, and a derivative order of 2. The wavenumber of key peaks was recorded. Spectra (without the Savitzky-Golay filter) were then exported to Fityk (version 1.3.1), smoothed with Voigt smoothing, and the peaks identified with the second derivative were added to the software. A new spectrum was obtained from the autofit function, and the fit was compared to the original spectrum. After the fit was confirmed, the height and area of the CH_2_ and CH_3_ peaks were recorded and used to calculate the R_3/2_.

## Acknowledgements

We thank the DCU Core Facilities team for technical support. The research leading to these results has received funding from the European Research Council (ERC) under the European Union’s Horizon Europe research and innovation programme (grant agreement No 1101114969) and from Research Ireland (Pathway award 22/PATH-S/10692).

