## Supplementary Information for "Protocells reproduce morphological and chemical signatures of early microfossils"

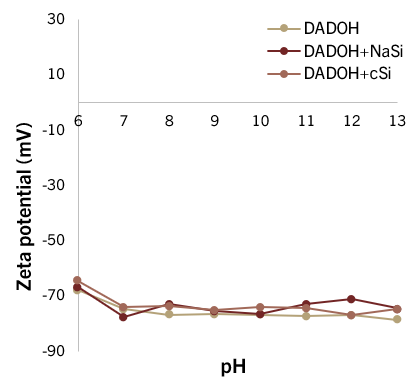

**Figure S1:** Surface charge of DADOH vesicles in solution, in control conditions or in the presence or Na_2_SiO_3_ or colloidal silica (300 ppm Si).

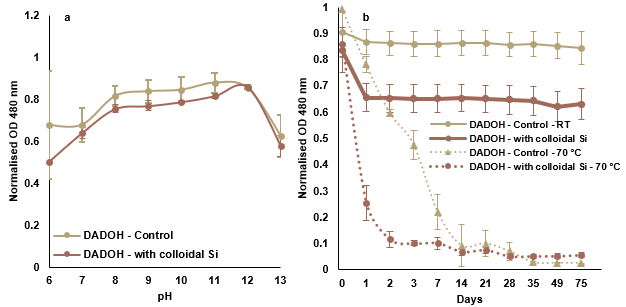

**Figure S2:** (a) Absorbance measurements at 480 nm of DADOH protocells across a pH range from 6 to 13, in control conditions (yellow) or exposed to colloidal Si (300 ppm Si) (dark orange). (b) Absorbance measurements at 480 nm, showing the stability over 8 weeks of DADOH protocells at pH 9, stored at room temperature (circles) or at 70 °C (triangles), in control conditions (yellow) or exposed to 300 ppm Si (dark orange).

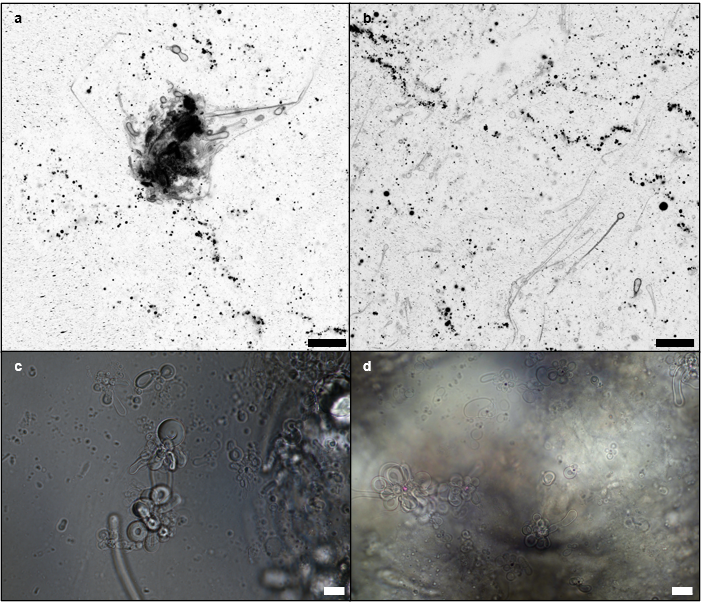

**Figure S3:** (a-b) CSLM micrographs showing the presence of vesicles in a highly concentrated SiO_2_ mixture, just before gelation. Protocells were stained with Rhodamine 6g. (c-d) Optical microscopy pictures showing the presence of protocells in the gel, a few minutes after gelation. Scale bars = 20 μm.

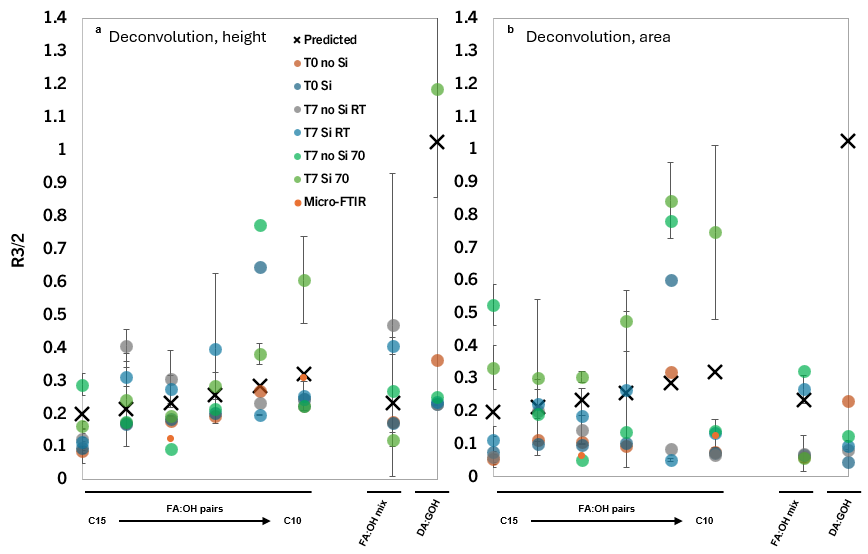

**Figure S4:** R_3/2_ values of protocells (C10 to C15 FA:OH pairs, FA:OH mix, and DA:GOH), in liquid solution or evaporated, after adding deconvolution to the spectra pre-processing. R_3/2_ values were obtained from (a) the height or (b) the area of the peaks of interest.
